# Twenty years of change in benthic communities across the Belizean Barrier Reef

**DOI:** 10.1101/2021.03.15.435443

**Authors:** Catherine Alves, Richard B. Aronson, Nadia Bood, Karl D. Castillo, Courtney Cox, Clare Fieseler, Zachary Locklear, Melanie McField, Laura Mudge, James Umbanhowar, Abel Valdivia, John F. Bruno

## Abstract

Disease, ocean warming, and pollution have caused catastrophic declines in the cover of living coral on reefs across the Caribbean. Subsequently, reef-building corals have been replaced by invertebrates and macroalgae, leading to changes in ecological functioning. We describe changes in benthic community composition and cover at 15 sites across the Belizean Barrier Reef (BBR) following numerous major disturbances—bleaching, storms, and disease outbreaks—over the 20-year period 1997–2016. We tested the role of potential drivers of change on coral reefs, including local human impacts and ocean temperature. From 1997 to 2016, mean coral cover significantly declined from 26.3% to 10.7%, while macroalgal cover significantly increased from 12.9% to 39.7%. We documented a significant decline over time of the reef-building corals *Orbicella* spp. and described a major shift in benthic composition between early sampling years (1997–2005) and later years (2009–2016). The covers of hard-coral taxa, including *Acropora* spp., *M. cavernosa, Orbicella* spp., and *Porites* spp., were negatively related to marine heatwave frequency. Only gorgonian cover was related, negatively, to our metric of the magnitude of local impacts (the Human Influence Index). Changes in benthic composition and cover were not associated with local protection or fishing. This result is concordant with studies throughout the Caribbean that have documented living coral decline and shifts in reef-community composition following disturbances, regardless of local fisheries restrictions. Our results suggest that benthic communities along the BBR have experienced disturbances that are beyond the capacity of the current management structure to mitigate. We recommend that managers devote greater resources and capacity to enforce and expand existing marine protected areas and that government, industry, and the public act to reduce global carbon emissions.

## INTRODUCTION

Coral reefs worldwide have experienced remarkable changes over the past 50 years, particularly the widespread declines of reef-building corals and large, predatory fishes (1–7). These changes have caused a reduction in or effective loss of essential ecological functions, including the provisioning of habitat for fisheries production and the maintenance of reef structure for shoreline protection (8,9). Given the substantial economic and cultural value of healthy reefs (10), this degradation is affecting coastal human communities that depend on reefs for food, income, and protection from storms.

Numerous factors are responsible for the well-documented degradation of Caribbean reefs. Acroporid corals, which dominated Caribbean reefs for millions of years, experienced 90–95% mortality due to white-band disease in the 1980s (11). This disease, likely exacerbated by ocean warming (12), coupled with increased frequency and intensity of hurricanes (13–15), reduced the habitat complexity, or rugosity, of Caribbean reefs (16). Several other disease syndromes have greatly reduced the cover of other coral taxa, including black-band disease, which primarily affects brain corals (17), yellow-band disease, which primarily affects *Orbicella* spp. (18), and, more recently, stony coral tissue loss disease, which affects numerous species, including *Dendrogyra cylindrus, Pseudodiploria strigosa, Meandrina meandrites, Eusmilia fastigiata, Siderastrea siderea* and *Diploria labyrinthiformis* (19). Coral bleaching and other manifestations of ocean warming, including increased disease severity, are primary causes of coral loss in the Caribbean (20–27). On local scales, increased sedimentation from coastal development affects coral reefs by increasing turbidity and smothering corals (28,29). Secondary drivers include factors that have increased the cover of fleshy macroalgae (seaweeds), including the death of scleractinian corals and the consequent opening of space and other resources (30), nutrient loading, and the loss of herbivores, particularly the sea urchin *Diadema antillarum* due to a regional disease outbreak (31), and herbivorous fishes due to fishing (32–37).

Despite the clear and well-documented changes to Caribbean reefs, there is ongoing disagreement about the causes of and best remedies for reef decline (20,38–41). The crux of the debate is about the relative importance of local causes—pollution, eutrophication, fishing, and consequent seaweed blooms—compared with regional-to-global causes such as ocean warming and acidification. Scientists, agencies, and organizations that view localized drivers as predominant generally argue for local mitigation, the primary recommendation being fisheries restrictions, such as within Marine Protected Areas (MPAs) (34,42–44). In contrast, the view that anthropogenic climate change has been a significant or predominant cause of reef decline leads to the conclusion that without rapid cuts in carbon emissions, local protections and other localized management actions, such as restoration, will ultimately fail (20,39,45).

The purpose of this study was to measure changes to benthic communities of the Belizean Barrier Reef (BBR) from 1997 to 2016 and to determine whether they were related to protection status, fishing, local human impacts, and ocean-temperatures anomalies (i.e., ocean heatwaves). We performed surveys of the coral reef benthos at 15 sites between 1997 and 2016 (46–48). We found that benthic-community composition changed substantially during this period, and that the observed loss of corals was negatively related to ocean heatwaves and largely unaffected by local impacts, fishing or protection status.

## MATERIALS AND METHODS

### Study area

Scientists have tracked reef community composition across Belize for over 50 years, mostly in short-term, longitudinal studies (e.g., 11,46,48–50). Belize has an extensive, 30-plus-year-old MPA network (46) and a history of frequent large-scale disturbances (Table 1). We surveyed fore-reef benthic communities at 15–18 m depth at 15 sites along the BBR during the summer months in 1997, 1999, 2005, 2009, and 2016 (Figure 1; Table S1). Due to logistical and resource constraints, only three of the 15 sites were surveyed every year: Bacalar Chico, Middle Caye, and Tacklebox (Table S1). Study sites were selected to maximize spatial heterogeneity and include a range of protections or management zones (5,47). These management protections included five sites within fully protected (FP) zones (otherwise known as “marine reserves”), where only non-extractive activities are permitted, three sites within general-use (GU) zones, where fishing is permitted with some gear restrictions (e.g., prohibitions on longlines, gillnets, and the use of spearguns and slings with SCUBA) and modest fishing limits (e.g., catch-size limits for queen conch and lobster), and seven sites in unprotected (NP) zones, where fishing is not restricted (46). Note that national seasonal closures for some species (e.g., Nassau grouper) and bans (e.g., on catching parrotfishes) applied to all three zones.

**Figure 1.**
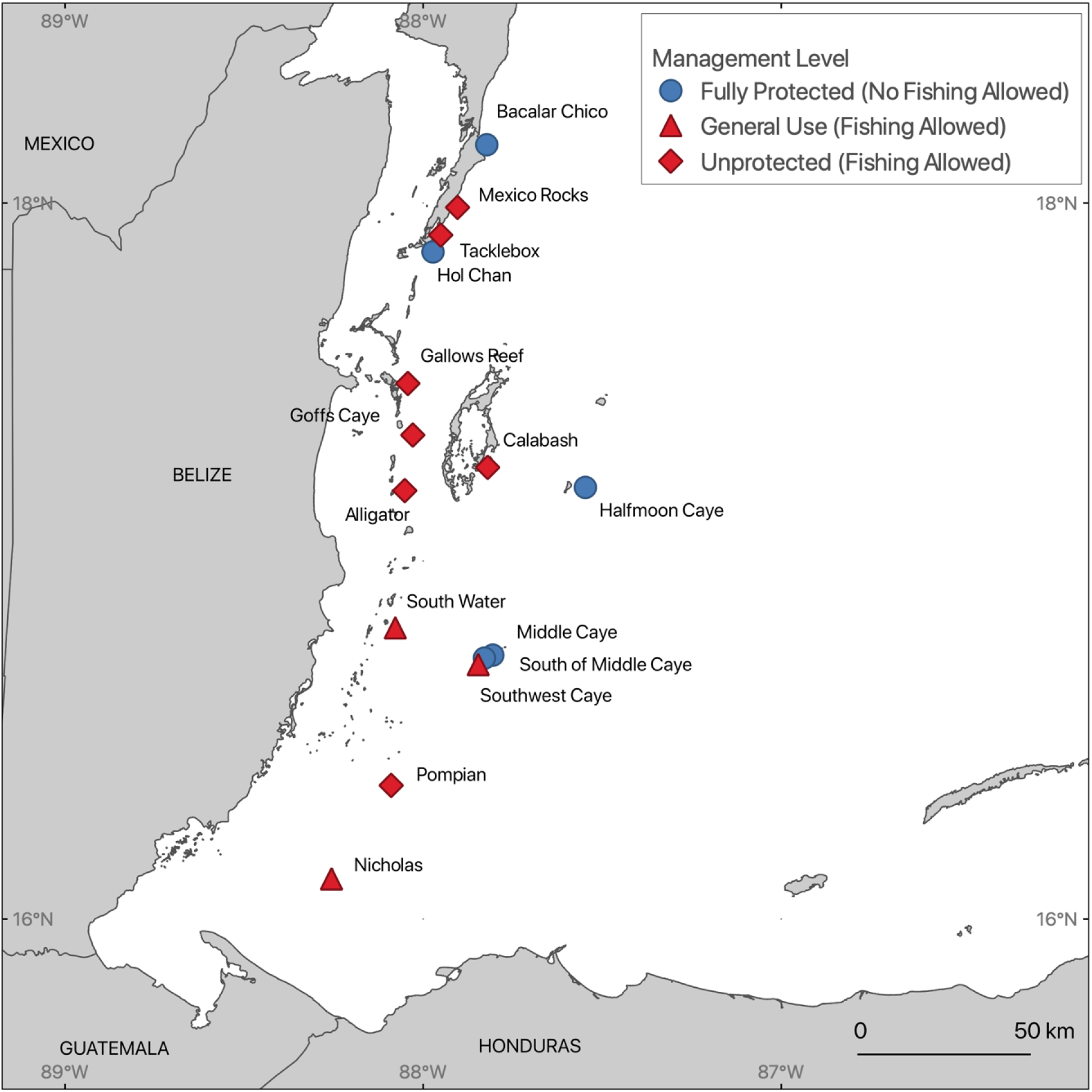
Study sites along the Belizean Barrier Reef. Sites are categorized by management and fishing level. Fishing is allowed in general use and unprotected sites (red), whereas fishing is prohibited in fully protected sites (blue).

**Table 1.** Timeline of major disturbances to the Belizean Barrier Reef.

| Year | Disturbance | References |
| --- | --- | --- |
| 1980s | Acroporid-specific white-band disease | (54) |
| 1983 | *Diadema*-specific disease | (31) |
| 1998 | Temperature-induced coral bleaching | (54,71) |
| 1998 | Hurricane Mitch | (94) |
| 2001 | Tropical Cyclone Iris | (95) |
| 2005 | Temperature-induced coral bleaching | (21,72–74) |
| 2007 | Hurricane Dean | (96) |
| 2009 | Earthquake | (97) |

### Benthic surveys

Benthic surveys were conducted *in situ* using SCUBA. At each site, dive teams laid out four to ten, 25-30 m x 2 m belt transects down the centers of reef spurs, perpendicular to the shoreline. The transects generally began on or near the shoulders of the spurs at 15–18 m depth, shoreward of the drop-off that characterizes most of the reefs, and ran upward toward the reef crest.

Transects were parallel to each other and were usually separated by > 10 m. Divers worked in buddy pairs, in which one diver laid out the transect tape and the other used a digital camera in an underwater housing to obtain videos and still-frame images of the benthos. At each site, we photographed or videotaped the belt transects at a standard distance of 25 cm above the benthos using a horizontal bar projected from the front of the camera housing. In all sampling years except 2016, we obtained underwater videos along the belt transects and extracted still frames from those videos (as outlined below). In 2016, we photographed the transects using a GoPro HERO4 by swimming at a rate of 5–7 minutes along the 30-m-long transect and taking a photograph every five seconds.

### Image extraction and analysis

Because of changes in imaging technology and analytical software over the course of this study we used several techniques to extract and analyze the benthic images from the underwater transects. For sampling year 1997, we recorded Hi-8 video of each transect, using two 30-watt ultrabright lights for illumination; in 1999 and 2005, we used Sony 3chip mini DVR without illumination. From these video cassettes we randomly selected 50 frames per transect, processed the images by de-interlacing, sharpening, and enhancing them, and saved them onto a CD-ROM. In 2009, we switched to digital video. We extracted the images from the video at a rate of 1-fps using Adobe Premiere Pro CC 2014. We ran the images through the Automator program in OS-X software to select every third, fifth or seventh image, depending on the length (in time) of the transect. We analyzed 15 images/transect/site for 2009 and 2016 because we found that we could obtain a similar level of inference about community composition with 15 images per transect as with the 50 images per transect suggested by Aronson et al. (51). To select the images, we automated the process using a code in R version 3.6.3 [1] to randomly choose, copy, and paste 15 images into a new folder from our source-folder of all images.

We analyzed the benthic cover of images from 1997–2005 using Coral Point Count software (52), and from 2009 and 2016 using CoralNet (53). We manually input species-level benthic identifications for each of 10 random points overlaid onto each image (51). When species-level identifications were not possible, benthic components were identified to genus or family. All benthic components identified were pooled into five benthic categories: (1) crustose–turf–bare space (abbreviated CTB), which represents substrate that is bare, dead, covered in turf algae, and/or crustose coralline algae (48,54), (2) hard corals (which includes all scleractinian corals and milleporine fire corals), (3) macroalgae, including algae in the genus *Halimeda,* (4) gorgonians, and (5) sponges. The corals *Orbicella annularis, O. favelota,* and *O. franksii* were pooled as *Orbicella* spp. because the species complex was not divided into three distinct species during the 1997 and 1999 data collection and because they were difficult to distinguish in some video frames. In all instances, image-level point-count data were converted to percent-cover estimates, and we calculated overall mean percent covers of each category for each site and year.

### Putative drivers of benthic community dynamics

We estimated local human impacts using the Global Human Influence Index (HII, version 2) from NASA’s Socioeconomic Data and Applications Center (SEDAC) database (55). The HII is a global dataset of 1-km grid cells aggregated from 1995–2004 designed to estimate location-specific human influence and thus potential impacts to natural populations and ecosystems via local direct and indirect human activities (e.g., harvesting and pollution). It is based on nine global data layers including human population density, land use, and access (which is estimated from coastlines, roads, railroads and navigable rivers). These aspects of human communities are known to be predictive of local human impacts in many natural systems including coral reefs (6,7,28,56–59). We extracted HII values for the BBR (Fig. S1) and calculated the sum of the HII scores of grid cells within a 50-km, 75-km, and 100-km buffer from the center-coordinates of each study site (Table S1). We used HII scores within the 50-km buffer for the final analysis because this metric performed well in exploratory models and it has been used successfully in prior work (5, 56). We then tested whether this index of local human impacts was related to observed changes on the monitored benthic reef communities.

Our measure of ocean-heatwave events was the site-specific frequency of Thermal Stress Anomalies (TSA Freq), obtained from the Coral Reef Temperature Anomaly Database (CoRTAD, Version 6) (60,61) (Fig. S2, Table S2). We used this metric to test for effects of thermal stress on the measured benthic groups. TSA Freq is defined as the number of deviations of 1 °C or greater from maximum weekly climatological sea-surface temperature during the 52 weeks preceding a reef survey. Other studies have found that TSA Freq is a significant predictor of coral-cover loss and coral-disease prevalence (62–64). The CoRTAD is based on 4-km-resolution weekly measurements made by the Advanced Very High-Resolution Radiometer (AVHRR) sensor (Pathfinder 5.0 and 5.2) beginning in 1982. Daytime and nighttime data were averaged weekly using data with a quality flag of 4 or better.

### Data analyses

To analyze changes in benthic composition and test for the effects of potential drivers of change, we built generalized linear mixed models (GLMM) in a Bayesian setting using the *blme* package (65). The response variables were the logit-transformed percent covers of key benthic categories. The final models had Year, Fishing level (“fishing”, which were the sites within FP zones, and “no fishing” which included GU and NP sites), HII at the 50-km buffer, and TSA Freq as fixed effects; and Site as a random effect. A blme prior with a *wishart* distribution was imposed over the covariance of the random effect and modeled coefficients. All predictor variables were additive, and the REML estimation was used to fit the data as it provides unbiased estimates for the variance components. Prior to fitting models, we rescaled and centered all numerical fixed effects to optimize comparisons among variables. The final model structure for each benthic category was as follows:

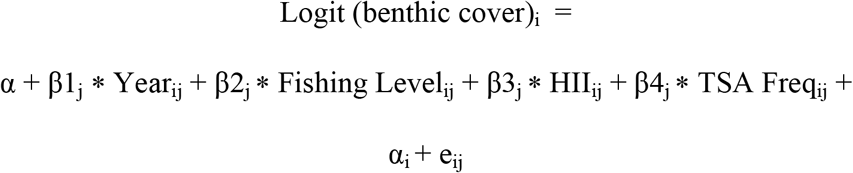

where α = intercept, α_i_ = random intercept (Site), e_jj_ = error term, and β_lj_ – β_4j_ are the coefficient estimates for covariates. The logit-transformed coral cover was modeled as an intercept (α), plus linear effects of Year, Fishing level, HII at 50km, and TSA Freq; a random intercept (α_i_) for Site, which is assumed to be normally distributed with a mean of 0 and variance σα^2^; and an error (e_ij_). The index *i* refers to sites (*i* = 1, …, 15), and *j* refers to the year of survey (*j* = 1997, …, 2016). The term e_jj_ was the within-site variance of benthic group cover and is assumed to be normally distributed with mean of 0 and a variance of σ^2^.

We evaluated collinearity among fixed factors by assessing variance-inflation factors and chose a threshold of 3 to determine correlated variables. We tested for homoscedasticity (equal variances across predictor variables) by plotting residuals against fitted values. Comparing fitted and residual values suggested that our models were reasonable models of the means. We also examined the marginal and conditional R-squared values of the models.

To examine changes in community composition of all benthic taxa within sites and across years, we constructed a non-metric multidimensional scaling (NMDS) ordination using the *vegan* package in R. We used the Bray–Curtis dissimilarity index to calculate distances among taxon-level cover data because it handles the large numbers of zeros (which denote absences) commonly found in ecological data and does not consider shared absences as being similar (66). To determine the effects of covariates (Year, TSA Freq, HII_50km, and Fishing level) on community composition changes of benthic taxa we ran a Permutational Multivariate Analysis of Variance (PERMANOVA) using the Bray-Curtis dissimilarity index to calculate distance matrices. All statistical analyses were performed in R version 3.6.3. The code and processed data are available at https://github.com/calves06/BRC.

## RESULTS

Among the five benthic groups of interest—hard corals, macroalgae, CTB, gorgonians, and sponges—we identified a significant decline in hard coral and CTB cover, significant increases in macroalgal and gorgonian cover, and no change in sponge cover (Figs. 2 & S3, Table 2). Fishing status (fished *versus* unfished) was not predictive of observed spatiotemporal variation in hard-coral, macroalgal, CTB, or sponge cover (Figs. 2 & 3, Table 2) and was marginally and negatively related to gorgonian cover. The Human Influence Index (HII) was also unrelated to hard-coral, macroalgal CTB, or sponge cover (Fig. 3, Table 2). HII was significantly and negatively related to gorgonian cover. TSA Freq, our metric of ocean-heatwave frequency, was significantly negatively related to the cover of hard corals and gorgonians, and unrelated to the cover of macroalgae, CTB, and sponges (Fig. 3).

**Figure 2.**
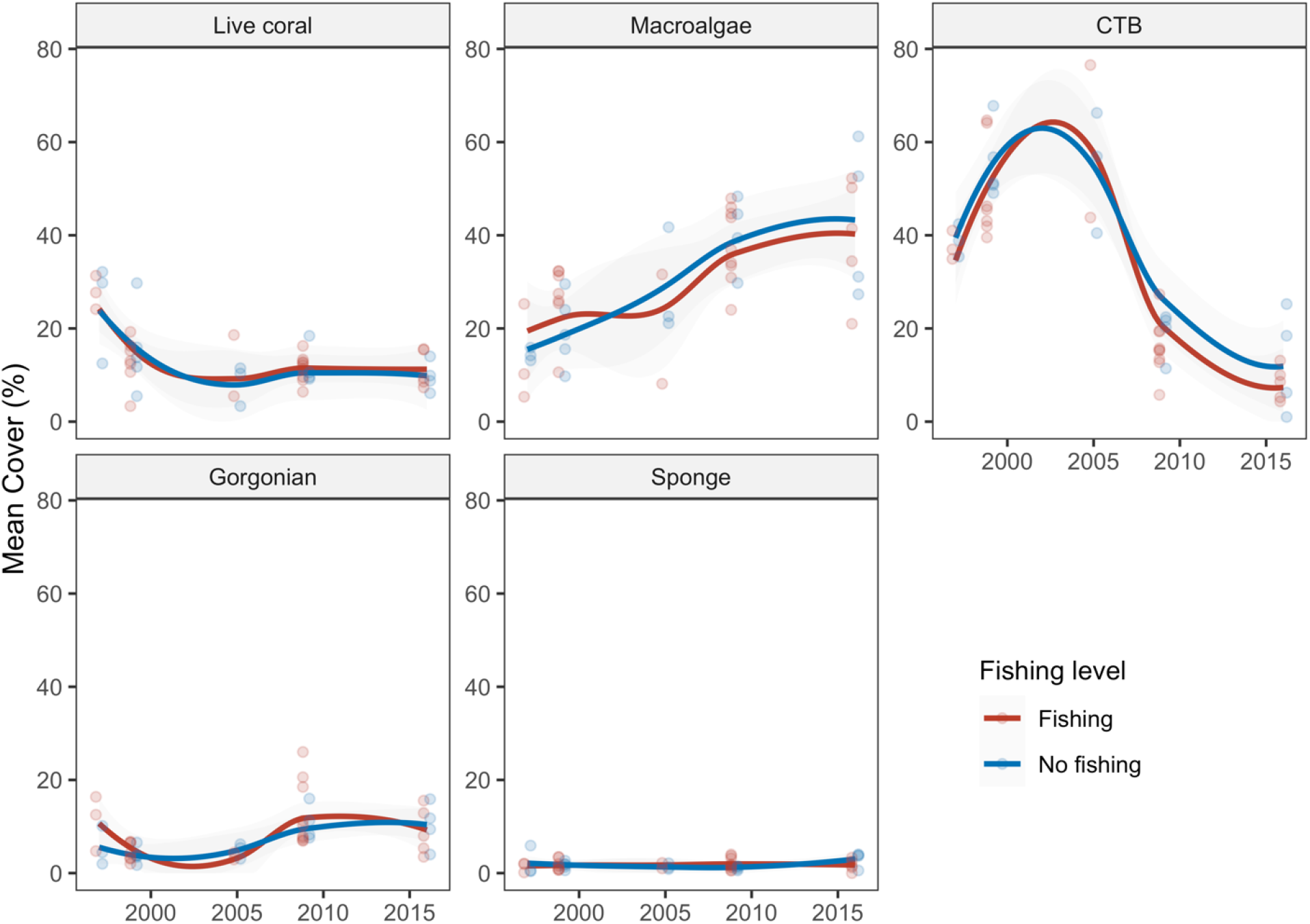
Percent cover of five benthic categories over time grouped by fishing level. Points are site means, lines are loess smoothed curves with a span of 1, shading indicates the 95% confidence intervals of the loess fits.

**Figure 3.**
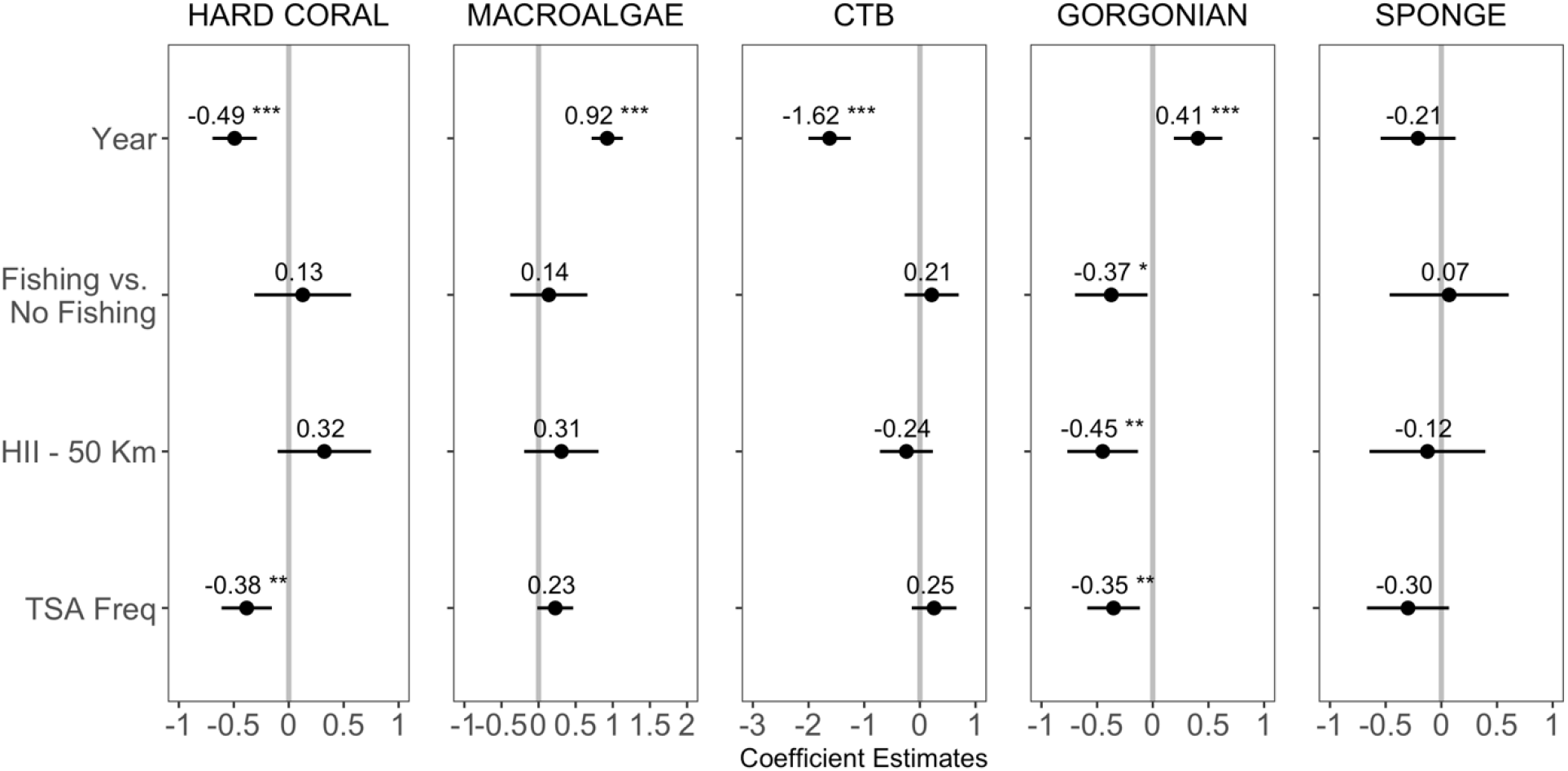
Effect-sizes (± 95% CI) of covariates from the Bayesian generalized linear mixed-effect model on the five benthic groups. Values above points are effect sizes. CIs crossing the vertical grey line represents a non-significant effect. Significance levels: *** = 0.001; ** = 0.01, * = 0.05.

**Figure 4.**
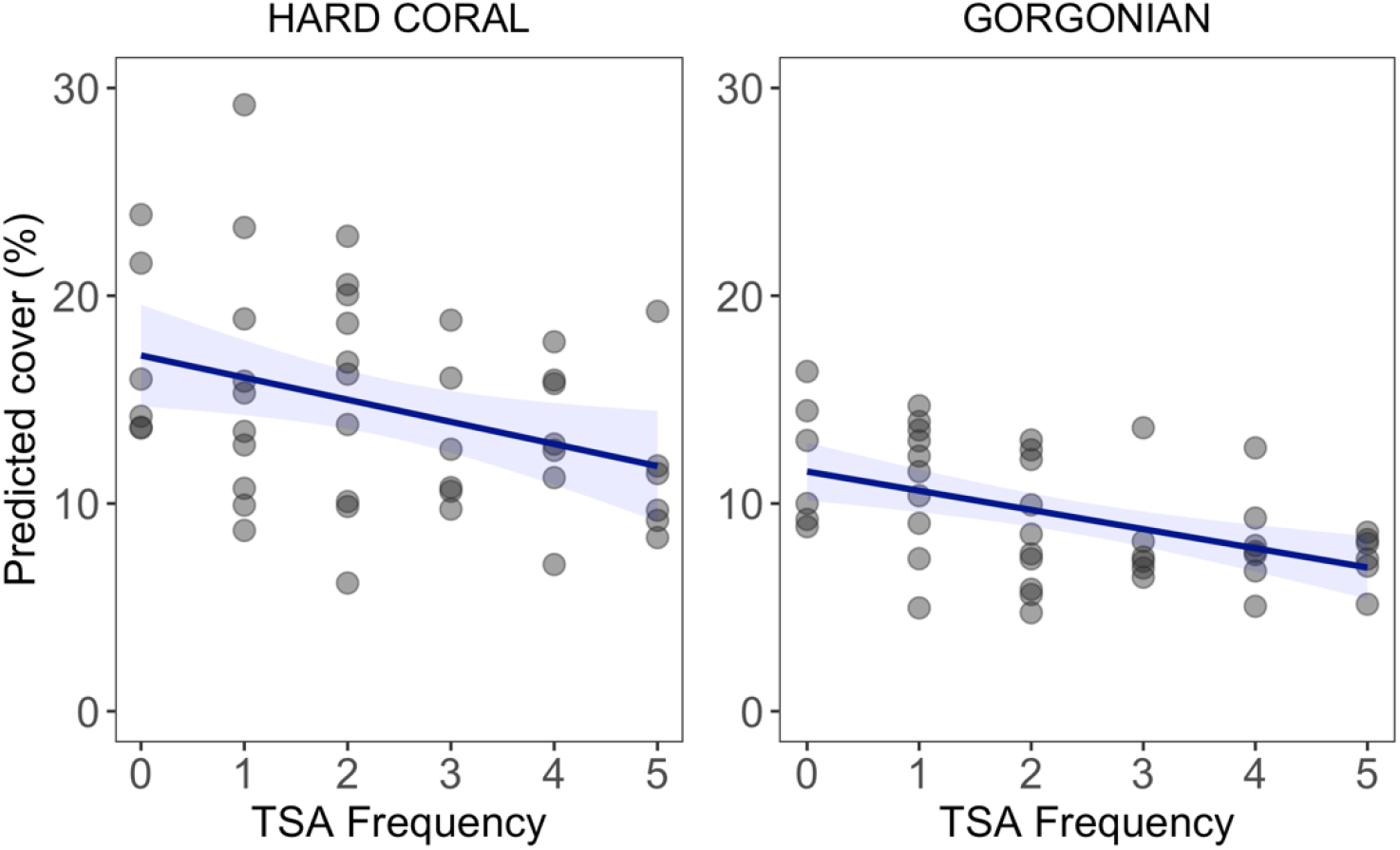
Predicted effect of TSA frequency on hard-coral and gorgonian cover. Points are predicted benthic group cover (back calculated from logit transformation) from Bayesian generalized liner mixed model accounting for time, fishing level, and human influence index. Blue lines are the fitted curves of the models and shaded areas are the 95 % CIs.

**Table 2.**
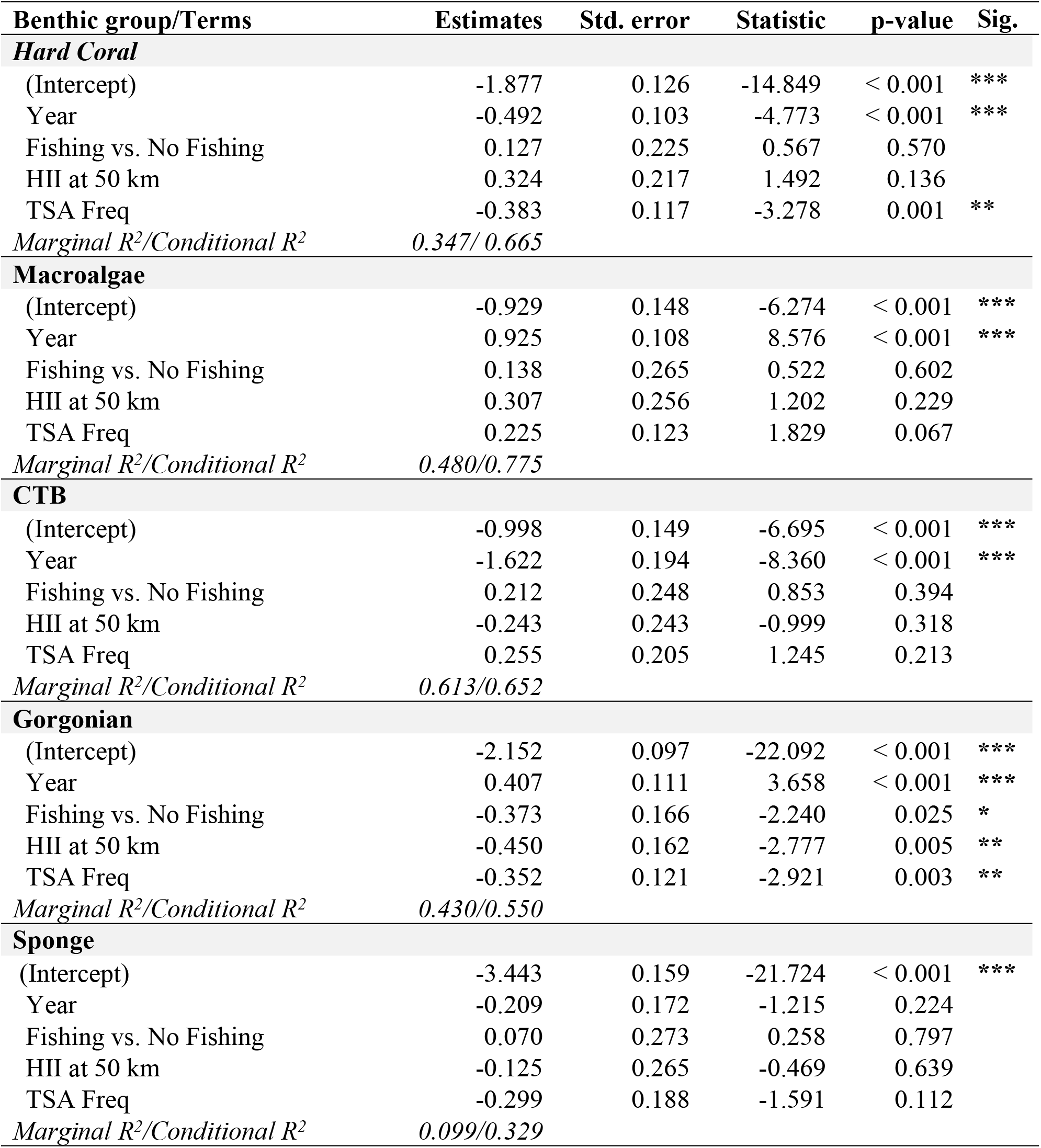
Estimated regression parameters for benthic groups coverage. Estimated regression parameters, standard errors, F-statistics, p-values, significance levels, and *marginal/conditional R^2^* from the final Bayesian generalized linear mixed models for each benthic group. Significance levels (Sig.) are: *** < 0.001; ** < 0.01, * < 0.05.

| Benthic group/Terms | Estimates | Std. error | Statistic | p-value | Sig. |
| --- | --- | --- | --- | --- | --- |
| **Hard Coral** |  |  |  |  |  |
| (Intercept) | -1.877 | 0.126 | -14.849 | < 0.001 | *** |
| Year | -0.492 | 0.103 | -4.773 | < 0.001 | *** |
| Fishing vs. No Fishing | 0.127 | 0.225 | 0.567 | 0.570 |  |
| HII at 50 km | 0.324 | 0.217 | 1.492 | 0.136 |  |
| TSA Freq | -0.383 | 0.117 | -3.278 | 0.001 | ** |
| Marginal R<sup>2</sup>/Conditional R<sup>2</sup> | 0.347/ 0.665 |  |  |  |  |
| **Macroalgae** |  |  |  |  |  |
| (Intercept) | -0.929 | 0.148 | -6.274 | < 0.001 | *** |
| Year | 0.925 | 0.108 | 8.576 | < 0.001 | *** |
| Fishing vs. No Fishing | 0.138 | 0.265 | 0.522 | 0.602 |  |
| HII at 50 km | 0.307 | 0.256 | 1.202 | 0.229 |  |
| TSA Freq | 0.225 | 0.123 | 1.829 | 0.067 |  |
| Marginal R<sup>2</sup>/Conditional R<sup>2</sup> | 0.480/0.775 |  |  |  |  |
| **CTB** |  |  |  |  |  |
| (Intercept) | -0.998 | 0.149 | -6.695 | < 0.001 | *** |
| Year | -1.622 | 0.194 | -8.360 | < 0.001 | *** |
| Fishing vs. No Fishing | 0.212 | 0.248 | 0.853 | 0.394 |  |
| HII at 50 km | -0.243 | 0.243 | -0.999 | 0.318 |  |
| TSA Freq | 0.255 | 0.205 | 1.245 | 0.213 |  |
| Marginal R<sup>2</sup>/Conditional R<sup>2</sup> | 0.613/0.652 |  |  |  |  |
| **Gorgonian** |  |  |  |  |  |
| (Intercept) | -2.152 | 0.097 | -22.092 | < 0.001 | *** |
| Year | 0.407 | 0.111 | 3.658 | < 0.001 | *** |
| Fishing vs. No Fishing | -0.373 | 0.166 | -2.240 | 0.025 | * |
| HII at 50 km | -0.450 | 0.162 | -2.777 | 0.005 | ** |
| TSA Freq | -0.352 | 0.121 | -2.921 | 0.003 | ** |
| Marginal R<sup>2</sup>/Conditional R<sup>2</sup> | 0.430/0.550 |  |  |  |  |
| **Sponge** |  |  |  |  |  |
| (Intercept) | -3.443 | 0.159 | -21.724 | < 0.001 | *** |
| Year | -0.209 | 0.172 | -1.215 | 0.224 |  |
| Fishing vs. No Fishing | 0.070 | 0.273 | 0.258 | 0.797 |  |
| HII at 50 km | -0.125 | 0.265 | -0.469 | 0.639 |  |
| TSA Freq | -0.299 | 0.188 | -1.591 | 0.112 |  |
| Marginal R<sup>2</sup>/Conditional R<sup>2</sup> | 0.099/0.329 |  |  |  |  |

Throughout the two decades of this study, the substantial decline in hard-coral cover across the Belizean Barrier Reef from 26.3 % (± 7.3 SD) to 10.6 % (± 3.5 SD) (Fig. 2) was driven by a handful of reef-building coral species (Fig. 5). Notably, there was a significant decline of *Orbicella* spp., with mean cover falling from 12.7 % (± 7.4 SD) in 1997 to 2.2 % (± 0.9 SD) in 2016 (Fig. 5, Table S4; model estimate = - 0.719, p < 0.001). This decline was predominantly observed from 1997 to 1999, which included a major bleaching event and Hurricane Mitch (Fig. 5, Table 1), and from 2005 to 2009, which included a second bleaching event, Hurricane Dean, and an earthquake (Fig. 5, Table 1). The cover of hard-coral taxa such as *Acropora* spp., *Colpophyllia natans,* and the combined cover of “other coral” taxa (e.g., *Mycetophyllia* spp., *Madracis* spp., *Favia* spp. see Table S5 for a complete list) also declined significantly from 1997 to 2016 (Fig. 5, Table S4). The cover of the coral taxa *Agaricia agaricities, Diploria/Pseudodiploria* spp., *Montastrea cavernosa, Siderastrea* spp., *Porites astreoides,* and *Porites* spp. (*P. porites*, *P. furcata, and P. divericata*) remained relatively low and did not change significantly during the study period (Fig. 5, Table S4). The cover of *Agaricia tenuifolia* slightly but significantly increased (Fig. 5, Table S4). Fishing level and HII were not significant predictors of spatial and temporal changes of any coral taxa (Table S4), except for *P. astreoides,* for which sites with higher cover were associated with areas of higher HII (Table S4). The cover of *Acropora* spp., *M. cavernosa, Orbicella* spp*. Porites* spp., and “other coral” taxa were negatively correlated with TSA frequency (Table S4).

**Figure 5.**
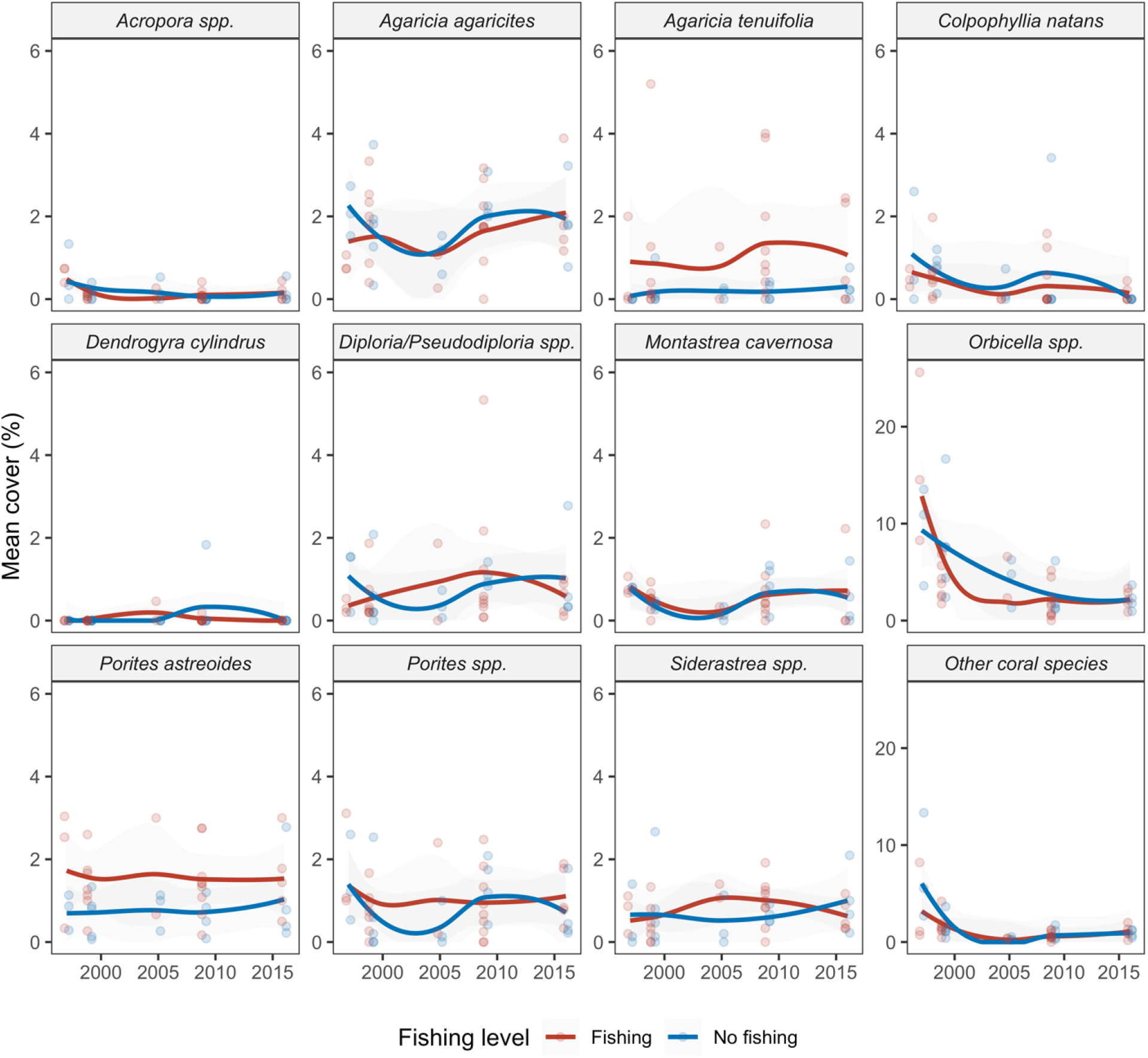
Mean percent cover of twelve taxonomic categories of hard corals, grouped by fishing level. Fishing occurs in red sites and is prohibited in blue sites. Points are site means for each surveyed year, lines are a loess smoothed curves with a span of 1, and shading indicates the 95% confidence intervals of the loess fits.

Based on the ordination analysis, there were major compositional shifts in the dominant benthic assemblages during 1997–2005 (left) and 2009–2016 (right) at every site (Fig. 6, Table 3), supporting the results of our models. The PERMANOVA showed that, among all covariates, time explained about 50% of the variability in benthic community changes (F = 45.8, p < 0.001) and was the only significant predictor of change in overall community composition (Fig. 6, Table 3). Fishing level, HII, and TSA frequency combined only accounted for 6% of community differences and were not good predictors of overall change of all taxa studied (Table 3). In 1997– 2005, the benthic communities of the BBR were dominated by CTB and long-lived, massive reef-building corals such as *Orbicella* spp. and *C. natans*. During 2009–2016, composition had shifted to domination by small and/or weedy hard-coral species, macroalgae, and gorgonians (Fig. 6).

**Figure 6.**
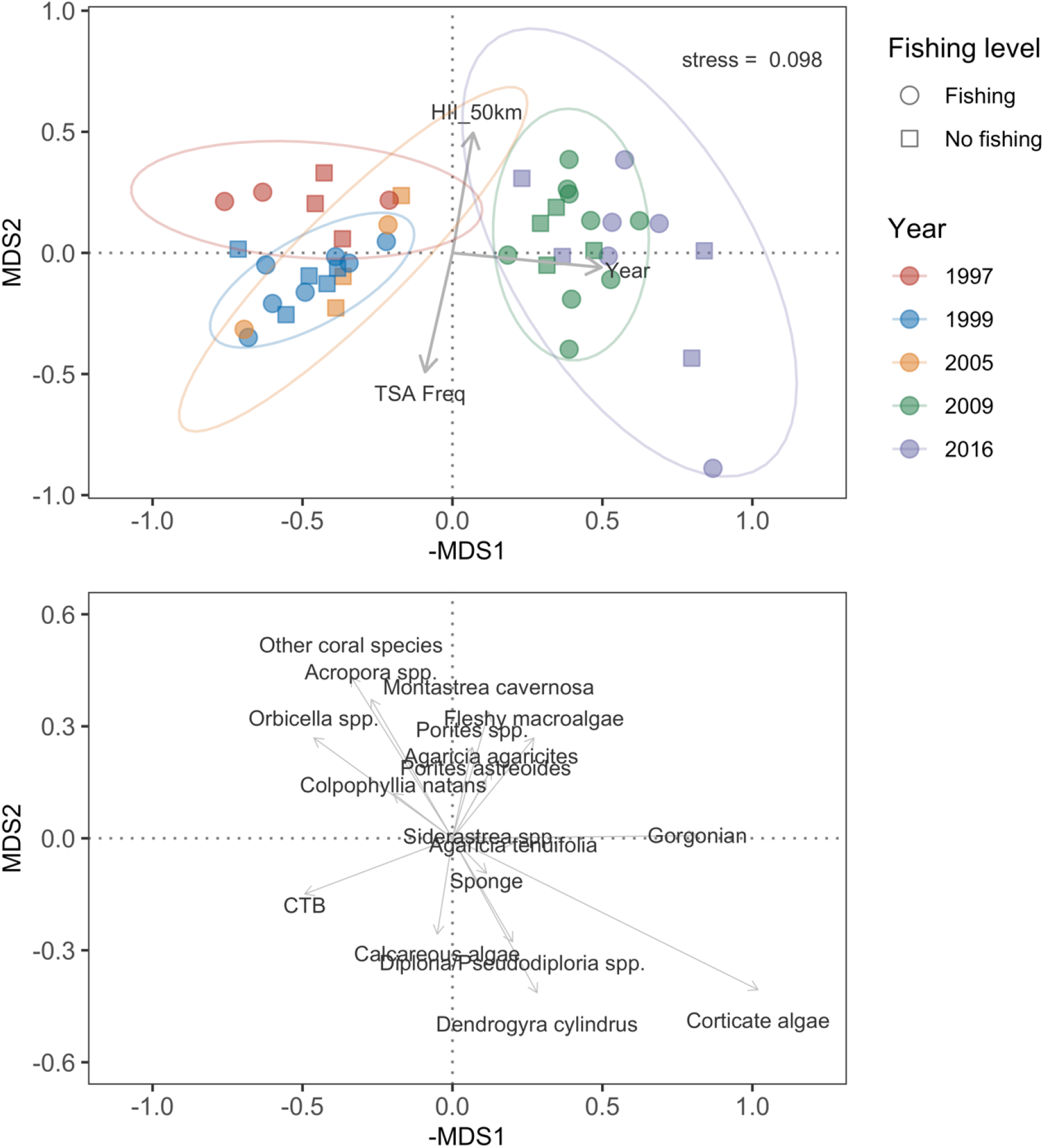
Non-metric multidimensional scaling (MDS) plot depicting taxon-level cover data colored by year. In the top panel, points represent individual sites, circles are fishing sites, and squares are no-fishing sites. Arrows represent the fitted loadings scores for Year, TSA_Freq, and HII_50km. In the bottom panel, the arrows and labels represent specific benthic categories loadings. The Bray–Curtis dissimilarity matrix was used and the stress value was 0.098.

**Table 3.** Results of the Permutational Multivariate Analysis of Variance (PERMANOVA) using the Bray-Curtis dissimilarity index to determine the effects of covariates in changes of benthic community composition cover. df: degrees of freedom, SS: sum of squares. Significance level (Sig.): *** < 0.001

| **Term** | **df** | **SS** | **R<sup>2</sup>** | **F** | **Pr(>F)** | **Sig.** |
| --- | --- | --- | --- | --- | --- | --- |
| Year | 1 | 2.347 | 0.501 | 45.761 | <0.001 | *** |
| HII 50km | 1 | 0.137 | 0.029 | 2.679 | 0.062 |  |
| TSA Freq | 1 | 0.072 | 0.015 | 1.401 | 0.218 |  |
| Fishing level | 1 | 0.075 | 0.016 | 1.465 | 0.200 |  |
| Residual | 40 | 2.051 | 0.438 |  |  |  |
| Total | 44 | 4.683 | 1.000 |  |  |  |

## DISCUSSION

Belize’s network of protected areas, designed and implemented in part to prevent the degradation of benthic reef assemblages on the BBR, has not achieved this goal. Our results complement previous findings for Belize reporting the failure of individual MPAs or the network overall to protect and restore populations of overharvested reef fishes (5,46,67,68), but see (68). We documented a statistically and ecologically significant decline in hard-coral cover, an increase in macroalgae and gorgonians, and a substantial decline of CTB, regardless of protection status (Fig. 2). Similar coral declines in isolated, well-protected, and seemingly “pristine” locations have been documented at many other sites globally (69,70).

We found that the benthic assemblages changed over time and were ecologically distinct between the earlier and later sampling intervals (1997–2005 and 2009–2016) (Fig. 6). For instance, the hard corals *Acropora* spp. and *Orbicella* spp. were more often present and more dominant (both had higher relative and absolute cover) in the early sampling years, as opposed to fleshy macroalgae and gorgonians, which dominated during later sampling years. In contrast, the cover of ‘weedy’ coral taxa such as *Porites* spp. and *Agaricia* spp. remained relatively consistent throughout the course of the study (Fig 5). The striking decline in *Orbicella* spp. (Fig. 5) was likely due to mortality from coral bleaching in 1998 (54,71) and 2005 (21,72–74), Hurricane Mitch in 1998, Hurricane Dean in 2007, and yellow-band disease in the early 2000s (Table 1).

Our results are concordant with previous studies in Belize that documented shifts in hard-coral and macroalgal cover <u>(75)</u>. For example, the patch reefs of Glovers Reef atoll had ~0% hard coral and 20% fleshy-macroalgal cover in 1970–1971 but phase-shifted to 20% hard coral and 80% macroalgal cover by 1996–1997 (75). This change was due to massive declines in the reef-building corals *Acropora cervicornis, A. palmata,* and *Orbicella* spp., and large increases in the cover and biomass of fleshy and corticated seaweeds including *Lobophora*, *Dictyota*, *Turbinaria*, and *Sargassum*. Prior to the beginning of our study, acroporid abundance had already declined across much of the BBR due to both hurricanes and white-band disease (11,68,76). Most remaining *A. cervicornis* and *A. palmata* was killed by high ocean temperatures during the 1998 mass-bleaching event (54,71). A longitudinal study of *A. palmata* along the Mexican portion of the Mesoamerican Barrier Reef also reported declines in acroporids, with *A. palmata* decreasing from 7.7% in 1985 to 2.9% in 2012 (76).

We attribute changes in the benthic assemblages of coral reefs along the BBR primarily to the large-scale disturbances to the system over the last several decades, including seven hurricanes and two mass-bleaching events caused by anthropogenic climate change (Table 1). We measured the potential effects of several putative drivers, including local human impacts estimated using the Human Influence Index (HII) and the frequency of ocean heatwaves (TSA Freq). Our results indicate that the local impacts had no measurable effect on hard-coral cover. HII was, however, significantly and negatively related to changes in gorgonians and positively associated with the cover of *Porites astreoides.* There is abundant evidence that local impacts, including pollution, fishing, and coastal land-use practices, can severely impact coral populations (28). Yet even when these stressors are clearly present, they are often overwhelmed by the effects of large-scale disturbances including ocean heatwaves and storms (20,39,57,75).

Shifts in the dominant benthic assemblages have been documented across the Caribbean, linked to regional disturbances such as herbivore declines, coral diseases, and mass-bleaching events (2,3,69,77,78). Across seven subregions in the Caribbean, Schutte et al. (2) found significant declines in hard-coral cover and increases in macroalgal cover from 1970–2005. Corals failed to recover in the Florida Keys (79) and the U.S. Virgin Islands (80) due to subsequent, repeated disturbances. The coral reefs of Bonaire exhibited similar trends over 15 years of bleaching, storms, and diseases, with a 22% decline in coral cover and an 18% increase in macroalgal cover by 2017 (81). These trends were also apparent in our study.

The primary management response to reef degradation has been implementation of MPAs (34,41,44,82). MPAs and MPA networks are areas where extractive activities are regulated via fishing closures or gear restrictions among others. Within well-designed and enforced MPAs, fish abundance and diversity often increase and in some cases spill over into adjacent, non-protected areas (68,83–86). Some MPAs also reduce other extractive activities that could directly or indirectly impact coral populations. However, a large majority of studies have found that MPAs are not slowing or preventing the decline of reef-building corals (50,63,67,79,87–89). A recent meta-analysis of 18 studies, encompassing 66 MPAs, reported that MPAs did not affect coral loss or recovery in response to large-scale disturbances including disease, bleaching, and storms (39). Our results for the BBR agree with this broad consensus.

Unlike local human impacts, anthropogenic climate change was clearly a significant driver of the dramatic shifts in community composition that occurred on the BBR over the two-decade study. Overall coral cover, and the cover of four coral taxa—*Acropora* spp. *Orbicella* spp., *Montastrea cavernosa,* and *Porites* spp.—were negatively related to heatwave frequency (Figs. 3 and 4). This result is in agreement with other studies that have documented coral mortality and consequent declines in coral cover following the temperature-induced mass-bleaching events on the BBR in 1998 and 2005 (48,54). Many other studies have documented the role of ocean heatwaves in coral decline around the world (21,22,63,69,88–93).

Our data show a substantial shift in the state of coral reefs along the Belizean Barrier Reef over a two-decade period rife with large-scale disturbances. The results illustrate the shortcoming of protected areas in mitigating these impacts. We documented declines in the key reef-building coral genera *Acropora* and *Orbicella*, subsequent increases in macroalgal and gorgonian cover, and an overall change in the benthic assemblages over the two-decade study. Ocean-heatwave frequency was the only significant predictor of coral population declines over time. Our results provide insight into the overriding influence of regional, and global drivers at a time of rapid climate change, which will help managers improve their decision-making.

## Acknowledgements

We thank the many volunteers who assisted with data collection, logistical support and image analysis over the years. We are extremely grateful for our partnerships with staff from the Belize Fisheries Department, Belize Coastal Zone Management Project, Wildlife Conservation Society, Marine Research Center at the University of California–Berkeley, Belize Audubon Society, Pelican Beach Resort, Rum Point Inn, Sea Sports Belize, Healthy Reefs for Healthy People, Bacalar Chico National Park and Marine Reserve, Hol Chan Marine Reserve, The Nature Conservancy, Southern Environmental Association, Toledo Institute for Development and Environment and the Smithsonian Institution.

## Data Availability

All relevant code and data are available at here: https://github.com/calves06/Belizean_Barrier_Reef_Change

## Permitting

The research was performed under permits from the Belize Fisheries Department to MM, NB, KC, CF, CC, and JFB including permit numbers 000018-09 and 000028-11.

## Author contributions

MM, RBA, NB, CF, and JFB designed the study

MM, RBA, NB, CC, CF, and JFB obtained the study funding

MM, NB, KC, CC, CF, CA, LM, AV, and JFB performed the surveys

MM, NB, CF, CC, and JFB performed the pre-survey expedition planning

CA, MM, NB, CF, CC, and AV organized the study data

MM, NB, CF, CA, ZL, LM, and JFB analyzed the benthic videos and images

CA, JU, AV, and JFB analyzed the data

CA, RA, AV, and JFB wrote the manuscript with contributions from the other authors

## LITERATURE CITED

1. Bruno JF, Selig ER. Regional decline of coral cover in the Indo-Pacific: timing, extent, and subregional comparisons. PLoS One. 2007;e711.

2. Schutte VGW, Selig ER, Bruno JF. Regional spatio-temporal trends in Caribbean coral reef benthic communities. Marine Ecology Progress Series. 2010;402:115–22.

3. Gardner TA, Côté IM, Gill JA, Grant A, Watkinson AR. Long-term region-wide declines in Caribbean corals. Science. 2003;301:958–60.

4. De’ath G, Fabricius KE, Sweatman H, Puotinen M. From the Cover: The 27-year decline of coral cover on the Great Barrier Reef and its causes. Proceedings of the National Academy of Sciences. 2012 Oct 1;109(44):17995–9.

5. Valdivia A, Cox CE, Bruno JF. Predatory fish depletion and recovery potential on Caribbean reefs. Sci Adv. 2017 Mar;3(3):e1601303.

6. Stallings CD. Fishery-independent data reveal negative effect of human population density on Caribbean predatory fish communities. PLoS ONE. 2009;4:e5333.

7. Ward-Paige CA, Mora C, Lotze HK, Pattengill-Semmens C, McClenachan L, Arias-Castro E, et al. Large-scale absence of sharks on reefs in the Greater-Caribbean: a footprint of human pressures. PLoS ONE. 2010;5:e11968.

8. Kuffner IB, Toth LT. A geological perspective on the degradation and conservation of western Atlantic coral reefs. Conservation Biology. 2016 Aug;30(4):706–15.

9. Perry CT, Murphy GN, Kench PS, Smithers SG, Edinger EN, Steneck RS, et al. Caribbean-wide decline in carbonate production threatens coral reef growth. Nat Commun. 2013 Jun;4(1):1402.

10. Spalding M, Burke L, Wood SA, Ashpole J, Hutchison J, zu Ermgassen P. Mapping the global value and distribution of coral reef tourism. Marine Policy. 2017 Aug;82:104–13.

11. Aronson RB, Precht WF. White-band disease and the changing face of Caribbean coral reefs. Hydrobiologia. 2001;460:25–38.

12. Randall CJ, van Woesik R. Contemporary white-band disease in the Caribbean has been driven by climate change. Nature Climate Change. 2015;5:375–9.

13. Webster PJ, Holland GJ, Curry JA, Chang HR. Atmospheric science: Changes in tropical cyclone number, duration, and intensity in a warming environment. Science. 2005;309(5742):1844–6.

14. Emanuel KA. Downscaling CMIP5 climate models shows increased tropical cyclone activity over the 21st century. Proceedings of the National Academy of Sciences of the United States of America. 2013;110(30):12219–24.

15. Elsner JB, Kossin JP, Jagger TH. The increasing intensity of the strongest tropical cyclones. Nature. 2008;455:92–5.

16. Alvarez-Filip L, Dulvy NK, Gill JA, Côté IM, Watkinson AR. Flattening of Caribbean coral reefs: region-wide declines in architectural complexity. Proceedings of the Royal Society B: Biological Sciences. 2009;276(1669):3019–25.

17. Edmunds PJ. Extent and effect of Black Band Disease on a Caribbean reef. Coral Reefs. 1991;

18. Bruckner AW, Bruckner RJ. Consequences of yellow band disease (YBD) on Montastraea annularis (species complex) populations on remote reefs off Mona Island, Puerto Rico. Diseases Aquatic Organisms. 2006;Special Issue 69:67–73.

19. Alvarez-Filip L, Estrada-Saldívar N, Pérez-Cervantes E, Molina-Hernández A, González- Barrios FJ. A rapid spread of the stony coral tissue loss disease outbreak in the Mexican Caribbean. PeerJ. 2019 Nov 26;7:e8069.

20. Aronson RB, Precht WF. Conservation, precaution, and Caribbean reefs. Coral Reefs. 2006;25:441–50.

21. Eakin CM, Morgan JA, Heron SF, Smith TB, Liu G, Alvarez-Filip L, et al. Caribbean corals in crisis: Record thermal stress, bleaching, and mortality in 2005. PLoS ONE. 2010;5(11).

22. Baker AC, Glynn PW, Riegl B. Climate change and coral reef bleaching: An ecological assessment of long-term impacts, recovery trends and future outlook. Estuarine, Coastal and Shelf Science. 2008 Dec;80(4):435–71.

23. Williams EH, Bunkley-Williams L. The world-wide coral reef bleaching cycle and related sources of coral mortality. Atoll Research Bulletin. 1990;335:1–71.

24. Manzello DP. Rapid Recent Warming of Coral Reefs in the Florida Keys. Sci Rep. 2015 Dec;5(1):16762.

25. Alemu I JB, Clement Y. Mass Coral Bleaching in 2010 in the Southern Caribbean. Dias JM, editor. PLoS ONE. 2014 Jan 6;9(1):e83829.

26. Winter A, Appledoorn RS, Bruckner A, Williams EH Jr, Goenaga C. Sea surface temperatures and coral reef bleaching off La Parguera, Puerto Rico (northeastern Caribbean Sea). Coral Reefs. 1998;17:377–82.

27. McWilliams JP, Côté IM, Gill JA, Sutherland WJ, Watkinson AR. Accelerating impacts of temperature-induced coral bleaching in the Caribbean. Ecology. 2005 Aug;86(8):2055–60.

28. Fabricius KE. Effects of terrestrial runoff on the ecology of corals and coral reefs: review and synthesis. Marine Pollution Bulletin. 2005;50:125–46.

29. Speare KE, Duran A, Miller MW, Burkepile DE. Sediment associated with algal turfs inhibits the settlement of two endangered coral species. Marine Pollution Bulletin. 2019 Jul;144:189–95.

30. Williams ID, Polunin NVC, Hendrick VJ. Limits to grazing by herbivorous fishes and the impact of low coral cover on macroalgal abundance on a coral reef in Belize. Marine Ecology Progress Series. 2001;222:187–96.

31. Carpenter RC. Mass mortality of Diadema antillarum. Marine Biology. 1990;104:67–77.

32. Hay ME, Colburn T, Downing D. Spatial and temporal patterns in herbivory on a Caribbean fringing-reef - the effects on plant-distribution. Oecologia. 1983;58:299–308.

33. Steneck RS, Mumby PJ, MacDonald C, Rasher DB, Stoyle G. Attenuating effects of ecosystem management on coral reefs. Sci Adv. 2018 May;4(5):eaao5493.

34. Mumby PJ, Harborne AR, Williams J, Kappel CV, Brumbaugh DR, Micheli F, et al. Trophic cascade facilitates coral recruitment in a marine reserve. Proc Natl Acad Sci USA. 2007;104:8362–7.

35. Paddack MJ, Reynolds JD, Aguilar C, Appeldoorn RS, Beets J, Burkett EW, et al. Recent Region-wide Declines in Caribbean Reef Fish Abundance. Current Biology. 2009 Apr;19(7):590–5.

36. Williams ID, Polunin NVC. Large-scale associations between macroalgal cover and grazer biomass on mid-depth reefs in the Caribbean. Coral Reefs. 2001;19:358–66.

37. Hughes T, Szmant AM, Steneck R, Carpenter R, Miller S. Algal blooms on coral reefs: What are the causes? Limnology and Oceanography. 1999;44:1583–6.

38. Aronson RB, Bruno JF, Precht WF, Glynn PW, Harvell CD, Kaufman L, et al. Causes of coral reef degradation. Science. 2003;302:1502–1502.

39. Bruno JF, Côté IM, Toth LT. Climate Change, Coral Loss, and the Curious Case of the Parrotfish Paradigm: Why Don’t Marine Protected Areas Improve Reef Resilience? 2018;30.

40. Pandolfi JM, Jackson JBC, Baron N, Bradbury RH, Guzman HM, Hughes TP, et al. Are U.S. coral reefs on the slippery slope to slime? Science. 2005;307:1725–6.

41. Russ GR, Questel S-LA, Rizzari JR, Alcala AC. The parrotfish–coral relationship: refuting the ubiquity of a prevailing paradigm. Marine Biology. 2015 Oct;162(10):2029–45.

42. Mumby PJ, Harborne AR. Marine reserves enhance the recovery of corals on Caribbean reefs. PLoS One. 2010;5(1):e8657.

43. Hughes TP, Bellwood DR, Folke C, Steneck RS, Wilson J. New paradigms for supporting the resilience of marine ecosystems. Trends In Ecology & Evolution. 2005;20:380–6.

44. West JM, Salm RV. Resistance and resilience to coral bleaching: Implications for coral reef conservation and management. Conservation Biology. 2003;17:956–67.

45. Bruno JF, Bates AE, Cacciapaglia C, Pike EP, Amstrup SC, van Hooidonk R, et al. Climate change threatens the world’s marine protected areas. Nature Clim Change. 2018 Jun;8(6):499–503.

46. Cox C, Valdivia A, McField M, Castillo K, Bruno JF. Establishment of marine protected areas alone does not restore coral reef communities in Belize. Marine Ecology Progress Series. 2017;563(1):65–79.

47. Bood N D. Recovery and resilience of coral assemblages on managed and unmanaged reefs in Belize: A long-term study. [Master’s Thesis]. [Mobile, AL]: University of South Alabama; 2006.

48. McField MD. Influence of disturbance on coral reef community structure in Belize. Proceedings 9’th International Coral Reef Symposium. 2000;1:63–8.

49. Stoddart DR. Post-hurricane changes on the British Honduras reefs: re-survey of 1972. Proceedings of the Second International Coral Reef Symposium. 1974;2:473–83.

50. McClanahan TR, McField M, Huitric M, Bergman K, Sala E, Nystrom M, et al. Responses of algae corals and fish to the reduction of macroalgae in fished and unfished patch reefs of Glovers Reef Atoll, Belize. Coral Reefs. 2001;19:367–79.

51. Aronson RB, Edmunds PJ, Precht WF, Swansop DW, Levitan DR. Large-scale, long-term monitoring of Caribbean coral reefs: simple, quick, inexpensive techniques. Atoll Research Bulletin. 1994;421.

52. Kohler KE, Gill SM. Coral Point Count with Excel extensions (CPCe): A Visual Basic program for the determination of coral and substrate coverage using random point count methodology. Computers & Geosciences. 2006 Nov;32(9):1259–69.

53. Beijbom O, Edmunds PJ, Roelfsema C, Smith J, Kline DI, Neal BP, et al. Towards Automated Annotation of Benthic Survey Images: Variability of Human Experts and Operational Modes of Automation. Chen CA, editor. PLoS ONE. 2015 Jul 8;10(7):e0130312.

54. Aronson RB, Precht WF, Toscano MA, Koltes KH. The 1998 bleaching event and its aftermath on a coral reef in Belize. Marine Biology. 2002;141(3):435–47.

55. WCS and CIESIN. Wildlife Conservation Society - WCS, and Center for International Earth Science Information Network - CIESIN - Columbia University (2005) Last of the Wild Project, Version 2, 2005 (LWP-2): Global Human Influence Index (HII) Dataset (Geographic). Palisades, NY: NASA Socioeconomic Data and Applications Center (SEDAC). https://doi.org/10.7927/H4BP00QC. Accessed 26 November 2018. In 2005.

56. Mora C. A clear human footprint in the coral reefs of the Caribbean. Proc R Soc B. 2008 Apr 7;275(1636):767–73.

57. Bruno JF, Valdivia A. Coral reef degradation is not correlated with local human population density. Sci Rep. 2016 Sep;6(1):29778.

58. Knowlton N, Jackson JBC. Shifting baselines, local impacts, and global change on coral reefs. PLoS Biology. 2008;6:e54.

59. Cinner JE, Maire E, Huchery C, MacNeil MA, Graham NAJ, Mora C, et al. Gravity of human impacts mediates coral reef conservation gains. Proc Natl Acad Sci USA. 2018 Jul 3;115(27):E6116–25.

60. Selig ER, Casey KS, Bruno JF. New insights into global patterns of ocean temperature anomalies: implications for coral reef health and management. Global Ecology and Biogeography. 2010;19:397–411.

61. Selig ER, Harvell CD, Bruno JF, Willis BL, Page CA, Casey KS, et al. Analyzing the relationship between ocean temperature anomalies and coral disease outbreaks at broad spatial scales. In: Coral reefs and climate change: science and management [Internet]. Washington, DC: American Geophysical Union; 2006. p. 111–28. Available from: internal- pdf://Seligetal2006-3775384832/Seligetal2006.pdf

62. Bruno JF, Selig ER, Casey KS, Page CA, Willis BL, Harvell CD, et al. Thermal stress and coral cover as drivers of coral disease outbreaks. PLoS Biology. 2007;5:e124.

63. Selig ER, Casey KS, Bruno JF. Temperature-driven coral decline: the role of marine protected areas. Global Change Biology. 2012 May;18(5):1561–70.

64. Maina J, McClanahan TR, Venus V, Ateweberhan M, Madin J. Global gradients of coral exposure to environmental stresses and implications for local management. PLoS ONE. 2011;6(8):14.

65. Chung Y, Rabe-Hesketh S, Dorie V, Gelman A, Liu J. A nondegenerate penalized likelihood estimator for variance parameters in multilevel models. Psychometrika. 2013 Oct 1;78(4):685–709.

66. Legendre P, Legendre L. Numerical Ecology. Third Edition. Amsterdam, The Netherlands: Elsevier; 2012.

67. Huntington BE, Karnauskas M, Lirman D. Corals fail to recover at a Caribbean marine reserve despite ten years of reserve designation. Coral Reefs. 2011 Dec;30(4):1077–85.

68. McClanahan T, Muthiga N. Change in fish and benthic communities in Belizean patch reefs in and outside of a marine reserve, across a parrotfish capture ban. Mar Ecol Prog Ser. 2020 Jul 9;645:25–40.

69. Sheppard C. Coral mass mortalities in the Chagos Archipelago over 40 years_ Regional species and assemblage extinctions and indications of positive feedbacks. Marine Pollution Bulletin. 2020;12.

70. Brainard RE, Oliver T, McPhaden MJ, Cohen A, Venegas R, Heenan A, et al. Ecological Impacts of the 2015/16 El Niño in the Central Equatorial Pacific. Bulletin of the American Meteorological Society. 2018 Jan 1;99(1):S21–6.

71. Aronson RB, Macintyre IG, Precht WF, Murdoch TJT, Wapnick CM. The expanding scale of species turnover events on coral reefs in Belize. Ecological Monographs. 2002;72(2):233–49.

72. Neal BP, Khen A, Treibitz T, Beijbom O, O’Connor G, Coffroth MA, et al. Caribbean massive corals not recovering from repeated thermal stress events during 2005–2013. Ecology and Evolution. 2017;7(5):1339–53.

73. Miller, J., Muller, E., Rogers, C., Waara, R., Atkinson, A., Whelan, K. R. T., Patterson, M., Witcher B. Coral diserase following massive bleaching in 2005 causes 60% decline in coral cover on reefs in the US Virgin Islands. 2009. p. 925–37.

74. Villamizar E, Díaz MC, Rützler K, De Nóbrega R. Biodiversity, ecological structure, and change in the sponge community of different geomorphological zones of the barrier fore reef at Carrie Bow Cay, Belize. Marine Ecology. 2014;35(4):425–35.

75. McClanahan TR, Muthiga NA. An ecological shift in a remote coral atoll of Belize over 25 years. 1998;(25):122–30.

76. Rodriguez-Martinez RE, Banaszak AT, McField MD. Assessment of Acropora palmata in the Mesoamerican Reef System. PLOS ONE. 2014;9(4):7.

77. Bruno JF, Sweatman H, Precht WF, Selig ER, Schutte VGW. Assessing evidence of phase shifts from coral to macroalgal dominance on coral reefs. Ecology. 2009;90:1478–84.

78. Bak RPM, Nieuwland G. Long-term change in coral communities along depth gradients over leeward reefs in the Netherlands Antilles. Bulletin of Marine Science. 1995;56:609–19.

79. Toth LT, van Woesik R, Murdoch TJT, Smith SR, Ogden JC, Precht WF, et al. Do no-take reserves benefit Florida’s corals? 14 years of change and stasis in the Florida Keys National Marine Sanctuary. Coral Reefs. 2014 Sep;33(3):565–77.

80. Edmunds PJ, Elahi R. The demographics of a 15-year decline in cover of the Caribbean reef coral Montastraea annularis. Ecological Monographs. 2007;77:3–18.

81. Steneck RS, Arnold SN, Boenish R, de León R, Mumby PJ, Rasher DB, et al. Managing Recovery Resilience in Coral Reefs Against Climate-Induced Bleaching and Hurricanes: A 15 Year Case Study From Bonaire, Dutch Caribbean. Front Mar Sci. 2019 Jun 7;6:265.

82. Hughes TP, Graham NAJ, Jackson JBC, Mumby PJ, Steneck RS. Rising to the challenge of sustaining coral reef resilience. Trends in Ecology & Evolution. 2010;25(11):633–42.

83. Russ GR. Do marine reserves export adult fish biomass? Evidence from Apo Island, central Philippines. Marine Ecology Progress Series. 1996;132:1–9.

84. Roberts CM, Bohnsack JA, Gell F, Hawkins JP, Goodridge R. Effects of marine reserves on adjacent fisheries. Science. 2001;294:1920–3.

85. Lester S, Halpern B. Biological responses in marine no-take reserves versus partially protected areas. Mar Ecol Prog Ser. 2008 Sep 11;367:49–56.

86. Chirico AAD, McClanahan TR, Eklöf JS. Community- and government-managed marine protected areas increase fish size, biomass and potential value. Bernardi G, editor. PLoS ONE. 2017 Aug 14;12(8):e0182342.

87. Coelho VR, Manfrino C. Coral community decline at a remote Caribbean island: marine no-take reserves are not enough. Aquatic Conservation: Marine and Freshwater Ecosystems. 2007 Nov;17(7):666–85.

88. Graham NAJ, McClanahan TR, MacNeil MA, Wilson SK, Polunin NVC, Jennings S, et al. Climate Warming, Marine Protected Areas and the Ocean-Scale Integrity of Coral Reef Ecosystems. PLoS ONE. 2008;3:e3039.

89. Darling ES, McClanahan TR, Côté IM. Combined effects of two stressors on Kenyan coral reefs are additive or antagonistic, not synergistic. Conservation Letters. 2010 Apr;3(2):122–30.

90. Glynn PW. Coral reef bleaching: ecological perspectives. Coral Reefs. 1993;12:1–17.

91. Brown BE. Coral bleaching: causes and consequences. Coral Reefs. 1997;16:S129–38.

92. Bruno JF, Siddon CE, Witman JD, Colin PL, Toscano MA. El Niño related coral bleaching in Palau, Western Caroline Islands. Coral Reefs. 2001;20:127–36.

93. Stuart-Smith RD, Brown CJ, Ceccarelli DM, Edgar GJ. Ecosystem restructuring along the Great Barrier Reef following mass coral bleaching. Nature. 2018 Aug;560(7716):92–6.

94. Jackson JBC, Donovan MK, Cramer KL, Lam VV. Status and Trends of Caribbean Coral Reefs: 1970-2012. IUCN; 2014.

95. Aronson RB, Macintyre IG, Precht WF. Event preservation in lagoonal reef systems. Geology. 2005;33(9):717–20.

96. Ibarra-García EC, Abarca-Arenas LG, Ortiz M, Rodríguez-Zaragoza FA. Impact of hurricane Dean on Chinchorro Bank coral reef (Western Caribbean): Temporal variation in the food web structure. Ecological Indicators. 2020 Nov;118:106712.

97. Aronson RB, Precht WF, Macintyre IG, Toth L. Catastrophe and the Lifespan of Coral Reefs. Ecology. 2012;93(2):110912084141001.

